# Voluntary wheel running provides pain relief but transiently exacerbates gait impairments in male and female mice with unilateral osteoarthritis

**DOI:** 10.64898/2026.02.27.708530

**Authors:** Roxana Florea, Sara Hestehave, Laura Andreoli, Annia Wright, Sandrine M Géranton

**Author notes:** corresponding author S.M.G.

## Abstract

**Objective:** Physical activity is a first-line therapeutic intervention for managing osteoarthritis-related pain and functional impairment. However, the growing literature questions the long-term relevance of exercise-induced improvements in patients, while pre-clinical research evidence base is limited by reliance on stressful, forced exercise paradigms which do not reflect voluntary engagement. Here, we aimed to investigate the effects of voluntary wheel running on the pain experience in mice with joint pain.

**Design:** We investigated the impact of free access to a running wheel on sensory, functional and affective outcomes following unilateral intra-articular injection of monoiodoacetate in single-housed male and female C57Bl/6J mice.

**Results:** Monoiodoacetate injection transiently reduced running activity in both sexes; however, females rapidly resumed and sustained high activity levels over a two-month period, while males showed a progressive decline in running distance. Active males and females showed improvements in the monoiodoacetate-induced hindpaw secondary mechanical hypersensitivity. Moreover, mechanical thresholds positively correlated with the distance ran after injury, suggesting a functional relationship between exercise and secondary pain relief. However, access to a wheel temporarily exacerbated several monoiodoacetate-induced gait impairments in both sexes. Finally, while there were no obvious effects of running on anxio-depressive-like behaviours or cognitive functioning, exercise significantly impacted stress-induced faecal output and phenotypic regulation of body weight.

**Conclusions:** Our findings suggest that persistent loading of an injured knee joint may compromise functional outcomes independently of pain relief away from the joint, underscoring a critical consideration for exercise-based therapeutic strategies in osteoarthritis.

## 1. Introduction

As one of the most common causes of chronic pain world-wide, osteoarthritis (OA) is characterised by an imbalance between repair and destruction mechanisms within the joints [1]. Despite the functional impairments associated with the progression of this condition, pain remains the defining clinical symptom that prompts patients to seek medical care. The limited short-term efficacy and adverse effects of current pharmacological interventions, combined with an incomplete understanding of OA pathophysiology, have prompted a shift in OA management toward prevention and early intervention. Accordingly, first line therapeutic strategies now emphasise patient education, weight management (where applicable) and physical activity [2,3].

Exercise interventions have been widely reported to improve joint function [4], alleviate pain [2] and reduce anxio-depressive symptoms [5], common comorbidities occurring in OA patients [6]. The analgesic effects of exercise are believed to result from many mechanisms, including the enhancement of joint functional integrity *via* regular mechanical loading, modulation of immune responses [7] and enhancement of endogenous analgesia [8]. However, a growing body of reports question the clinical relevance of exercise interventions and point to the modest effect sizes, short-lived improvements and methodological concerns across the literature [9–12], leaving exercise interventions for OA pain management a subject of debate.

In laboratory settings, aerobic exercise such as wheel running, swimming and treadmill running are widely used to investigate the impact of and mechanisms underlying exercise-induced analgesia. Improvements in pain-like behaviour and associated affective anxio-depressive behaviours have been extensively documented in animal models of short- and long-term pain states [13,14]. In OA models, exercise interventions have also been found to somewhat improve pain-like behaviours [15,16] and joint integrity [17,18] but not universally [19–21]. Concerns have been particularly raised regarding the translational validity of forced exercise paradigms, such as treadmill running and swimming, which are increasingly showed to introduce stress confounds [22,23]. Moreover, the impact of exercise has been understudied in female rodents, despite the higher incidence of OA among women [1].

Here, we examined the therapeutic potential of voluntary wheel running in singly-housed male and female mice that received a unilateral injection of monoiodoacetate (MIA) in the knee synovial space. Behavioural assessment over the subsequent two months revealed an improvement in hindpaw secondary hyperalgesia in both sexes, with no significant changes in the anxio-depressive-like behaviours, nor novelty preference seeking, despite a significant impact on stress-induced faecal output and exercise-driven regulation of body weight. Notably, voluntary exercise transiently exacerbated several MIA-induced alterations in gait characteristics compared with mice that had no access to wheel. These results suggest that sustained use of an injured knee joint can impair functional outcomes, even when pain at distant sites is attenuated, an important consideration for therapeutic recommendations regarding exercise in OA.

## 2. Methods

Extended methods and materials can be found in the supplementary file. See ***fig. S1*** for timeline of experimental procedures and behavioural tests carried out.

### 2.1 Animals and study design

A total of 16 male and 16 female C57Bl/6J mice, ordered from Charles River UK at 6 weeks of age, were single housed in individually ventilated cages containing regular cage enrichment, with the intervention cage also containing a vertical running wheel. Food and water were provided *ad libitum*, with animals being housed in an alternating 12/12h light/dark cycle, and temperature and relative humidity-controlled conditions (22 ± 2ºC, 50 ± 10%). All procedures were carried out in accordance with the guidelines of the UK Animals (Scientific Procedures) Act 1989 and subsequent amendments. ARRIVE guidelines were followed throughout. A minimum period of two weeks of habituation was allowed prior to MIA injection to ensure mice with access to the wheel would reach relatively stable levels of activity.

### 2.2 Activity recording

Running activity was quantified using an activity wheel monitoring system (Columbus Instruments, Columbus, OH, USA). Intervention cages were equipped with a vertical polycarbonate running wheel and wheel revolutions were automatically detected using a Hall-effect sensor connected to an eight-channel counter and Quad CI-Bus interface. Data were recorded using Multi-Device Interface (MDI) software (version 1.11.7). Running activity was continuously monitored over a 24h period.

### 2.3 Monoiodoacetate-induced arthritis

To induce osteoarthritis, mice received an intra-articular injection of monoiodoacetate (MIA) in the left knee joint as previously reported [24]. On the day of injection, mice were placed under general anaesthesia and 10μl of 10% w/v MIA solution (Sigma-Aldrich) was injected in the synovial space.

### 2.4 Mechanical sensitivity

Hindpaw mechanical sensitivity was assessed using the up-and-down method as previously described [24]. Mice were individually placed in plexiglass cubicles situated on an elevated wire grid (Ugo Basile). Calibrated von Frey monofilaments (Ugo Basile, Italy) were applied to the plantar surface of the hindpaw to determine the 50% withdrawal threshold, starting with a monofilament of 0.6g, and using the formula: 50% *mechanical threshold* (log(*g*)) = 10^log(last filament weight (g))+*k*\*0.3^, where *k* is a constant value corresponding to each individual response pattern obtained [25].

### 2.5 CatWalk gait analysis

Gait behaviour was analysed using the CatWalk^®^ XT 10.0 system (Noldus Information Technology), as we previously described [24]. A minimum of two compliant runs were included in the analysis, and the parameters analysed were based on previous literature on arthritis models [26].

### 2.6 Open Field test

Anxiety-like behaviour was assessed using a square open field arena. After room habituation, mice were placed individually in the center of the open field arena. Subject’s movements were live tracked and recoded for 5min using the Ethovision XT14 software (Noldus Information Technology).

Raw movement data (x-y coordinates) were exported and analysed using a custom Python script that divided the arena into three concentric circular zones: ‘Near Center’ zone (innermost region), ‘Away from Center’ zone (intermediate zone), and ‘Far Away from Center’ zone (border zone). The animal’s position as well as the distance travelled every 0.2s were used to compute the proportion of time spent and total distance travelled within each zone, respectively.

### 2.7 Sucrose Preference test

Anhedonia was assessed using the sucrose preference test as previously described [24]. Testing was conducted in the home cages over four consecutive days: two days to account for side preference, and another two to account for potential competing rewarding effects between running activity and sucrose consumption. Preference for 1% sucrose solution over regular water was quantified during the animals’ active phase.

### 2.8 Novel Object Recognition test

To quantify novelty preference, the novel object recognition (NOR) test was employed as previously described [24]. The test consisted of a 10min habituation phase, during which mice were exposed to two identical objects, followed 3h later by a 5min test phase during which one of the familiar objects was replaced with a novel one. Mouse movements were recorded using Ethovision XT14 software (Noldus Information Technology), and exploration time was calculated using the formula: *Exploration time* 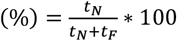 100, where *t*_*N*_ and *t*_*F*_ is the time spent exploring the novel and familiar object, respectively.

### 2.9 Faecal pellet output

To indirectly assess the stress responses mounted during von Frey testing sessions, the total number of faecal pellets produced by each group (active *vs* sedentary) throughout the testing period was recorded. The total pellet count was divided by the number of mice in each group, and the resulting values were plotted over time.

### 2.10 Statistics (see extended paragraph in supplementary information)

Data were analysed using GraphPad Prism (version 10.6.1 (892)) and IBM SPSS Statistics (version 30.0.0.0 (171)). Statistical analysis was mostly performed using Student’s *t* test, 1-way or 2-way repeated measures (RM) ANOVA, as well as simple linear regression. Normality was assessed using the Shapiro–Wilk test and inspection of Q–Q plots, which provide a more robust assessment when formal tests are overly sensitive. When sphericity or multivariate tests could not be computed due to an unbalanced data structure (*i*.*e*., an insufficient number of complete cases relative to the number of time points), we used linear mixed-effects models. For von Frey data, raw data were log-transformed to normalize distributions and stabilize variance. Where deviations from normality occurred (only in isolated subgroups), we used non-parametric tests (*e*.*g*. Mann–Whitney’s test). Significance was set at *P*<0.05. Data are presented as group means ± standard error of the mean (SEM), with the exception of box-and-whiskers plots, which display the IQR. Outliers were identified and excluded using Tukey’s method.

## 3. Results

### 3.1 Voluntary wheel running attenuates MIA-induced mechanical allodynia, but transiently exacerbates gait impairments

Following the intra-articular injection of MIA, both male (***Fig. 1A, B***: RM 1-way ANOVA, F_2, 14_=12.45, *P* = 0.0084) and female mice (***Fig. 1C, D***: mixed-effects 1-way ANOVA, F_2,14_=7.28, *P* = 0.0108) exhibited a reduction in voluntary running activity (phase 1). This phase of reduced activity was longer lasting in males, who ran significantly shorter distances until day 11 post-injection, compared to females which returned to baseline running by day 8. Both cohorts exhibited a rebound in activity (start of phase 2). This recovery was sustained in females, who maintained pre-injury activity levels for the remaining of the experiment. In contrast, the running distance in male mice continuously decreased from day 13 onwards. We observed similar, injury-independent sex differences using non-invasive recordings of home cage activity [27] (***fig. S2***).

**Figure 1.**
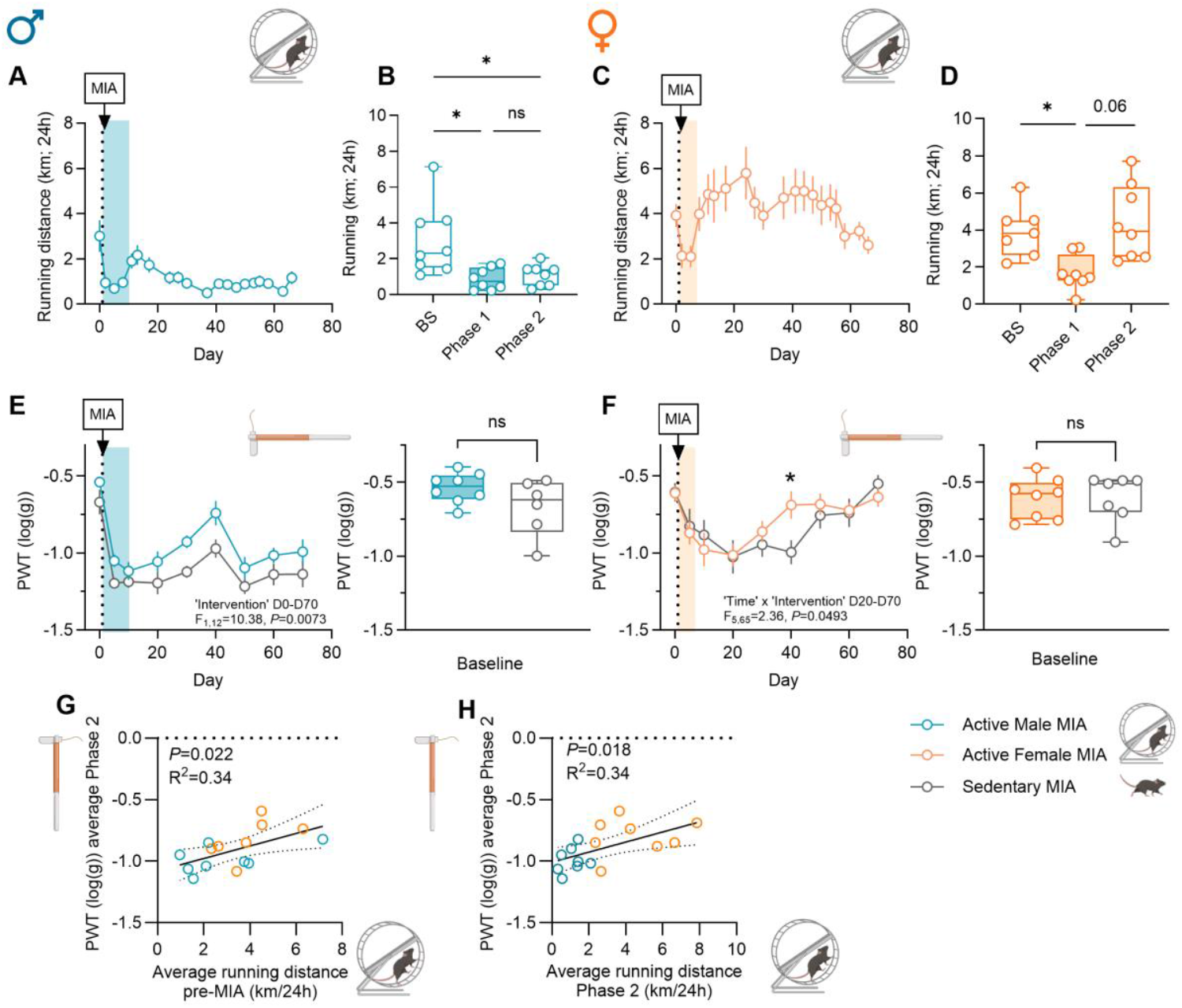
Wheel running attenuates MIA-induced mechanical allodynia across sexes. (**A**) Time course of average daily running distance and (**B**) wheel running activity in male mice separated into phases pre- and post-MIA; BS: baseline; Phase 1: day 2 to day 8; Phase 2: day 11 to end of experiment. \**P* < 0.05; *N* = 8. (**C**) Time course of daily running distance before and after the MIA injection and (**D**) wheel running activity in female mice separated into phases pre- and post-MIA; BS: baseline; Phase 1: day 2 to day 5; Phase 2: day 8 to end of experiment. \**P* < 0.05; *N* = 8. (**A, C**) Baseline measurements represent mean distance ran per day (24h) over the 7-day time period before MIA injection, while subsequent data points represent 3-5 day averages. (**E, F**) Mechanical allodynia at the ipsilateral hindpaw in male (**E**, *N* = 8/6) and female (**F**, *N* = 7/8) mice. (**G**) Correlation between average distance run pre-MIA (km/24h) and PWT ipsilateral (average Phase 2). (**H**) Correlation between average run Phase 2 (km/24h) and PWT ipsilateral (average Phase 2). \**P* < 0.05, \*\**P* < 0.01. (**A, C, E, F**) Coloured band indicates Phase 1. Data presented as mean ± SEM.

When looking at the pain-like behaviour, the knee MIA injection induced the development of robust mechanical allodynia at the ipsilateral hindpaw in both male (***Fig. 1E***, 2-way RM ANOVA, day 0 to 70: ‘Time’ F_8,96_=20.5, *P* < 0.0001) and female (***Fig. 1F***, ‘Time’ F_8,104_=9.06, *P* < 0.0001) mice, independently of wheel access. While there was no difference in baseline threshold due to the presence of the wheel (***Fig. 1E, F***, right panels), a significant effect of the intervention was observed in the male cohort across the time course of the experiment (***Fig. 1E***, 2-way RM ANOVA, day 0 to 70: ‘Intervention’ F_1,12_=10.38, *P* = 0.0073). Using a different set-up, we confirmed the analgesic effect of voluntary wheel running in males, while also observing enhanced hippocampal neurogenesis, a common consequence of exercise (***fig. S3***) [28]. The impact of running was not as obvious in females, despite their sustained increase in voluntary activity. However, females that had access to wheels did recover quicker than their controls, as evidenced by an interaction between time and intervention (***Fig. 1F***, 2-way RM ANOVA, ‘Time x Intervention’, day 20 to 70: F_5,65_=2.36, *P* = 0.049). When we looked at the relationship between the activity levels and mechanical thresholds across sexes (average Phase 2), we found a positive correlation between hindpaw sensitivity and the running activity both before and after the injection of MIA, suggesting a functional relationship between exercise and secondary pain relief (***Fig. 1G, H***).

Since both osteoarthritis-induced joint pain and functional deficits are associated with gait alterations, we also quantified gait performance using the CatWalk system. The MIA injection induced transient impairments in gait coordination and balance, impairments that peaked around day six post knee injection in both sexes (***Fig. 2A-I***). Wheel running exacerbated MIA-induced deficits, including those observed for the amount of time the injured hindpaw was in contact with the runway (single stance) and the amount of time the injured hindpaw was in ground contact during a gait cycle (duty cycle) in both males (***Fig. 2A***, D20-58: mixed-effects 2-way ANOVA, ‘Intervention’ F_1,12_=5.15, *P* = 0.043; ***B*:** D20-58: mixed-effects 2-way RM ANOVA, ‘Intervention’ F_1,12_=5.26, *P* = 0.041), and females (***Fig. 2C***, D12-67: mixed-effects 2-way RM ANOVA, ‘Intervention’ F_1,14_=8.48, *P* = 0.0045; ***D***: mixed-effects 2-way RM ANOVA, ‘Intervention’ F_1,14_=7.554, *P* = 0.0157). However, this was not the case for all temporal and coordination parameters analysed, as wheel running had no obvious impact on print area and maximum paw intensity during runway traversal (***Fig. 2E-H***).

**Figure 2.**
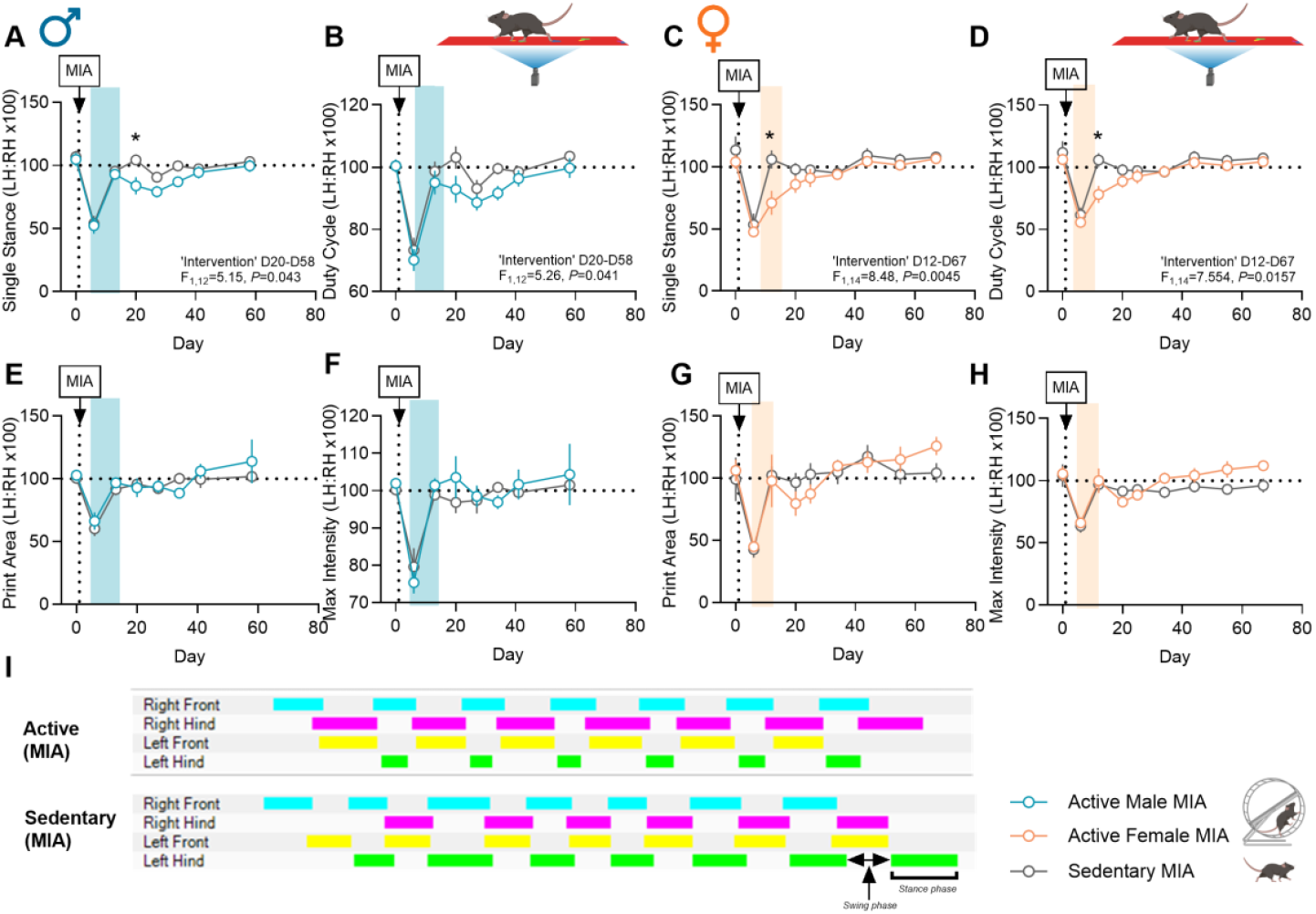
Wheel running transiently exacerbates gait impairments across sexes. (**A-D**) Time course of gait parameters impacted by access to a running wheel in male (**A, B;** *N* = 5-8) and female mice (**C, D**, *N* = 4-8). \**P* < 0.05, \*\**P* < 0.01. (**E**-**H**) Time course of gait parameters unchanged by access to a running wheel in male (**E, F**, *N* = 5-8) and female mice (**G, H**, *N* = 4-8). (**I**) Representative footprint timing view at day 12 after the MIA injection in female mice. (A-H) Coloured band indicates Phase 1. Data presented as mean ± SEM.

Together, these findings indicate that increased activity may have prolonged the negative consequences of MIA on joint function, as evidenced by exacerbation of gait alterations, while having some benefit on secondary mechanical sensitivity.

### 3.2 Voluntary wheel running does not obviously impact affective behaviours in male and female mice with unilateral OA but influences body weight gain

Persistent pain is known to promote the development of emotional and cognitive comorbidities, which not only decrease the quality of life, but can further exacerbate the pain symptoms [29,30]. Evaluation of affective and cognitive functioning was performed at two time points after the MIA injection, corresponding to periods associated with significant histopathological changes in the knee joint [31,32] and our previous observations of affective changes [27].

Depressive-like behaviour was evaluated using the sucrose preference test at one- and two-months post MIA injection. Because the rewarding effects of wheel running could compete with the rewarding effects of sucrose consumption [33,34], the test was run twice at each time point, once in the presence and once in the absence of the wheel for the active mice. At one month, voluntary wheel running did not significantly change sucrose preference in either male (***Fig. 3A***) or female (***Fig. 3B***) mice, irrespective of the presence of the wheel. However, we found a significant effect of the presence of the wheel on sucrose preference in males (***Fig. 3A***: RM 2-way ANOVA: ‘Wheel presence during sucrose test’ x ‘Intervention’: F_1,11_=5.097, *P* = 0.045), but not in females. Moreover, there was a significant correlation between the distance run and sucrose consumption when the wheels were removed (***fig. S4***), with preference for sucrose positively correlated with running. No significant differences were observed at two months (***Fig. 3C-D***).

**Figure 3.**
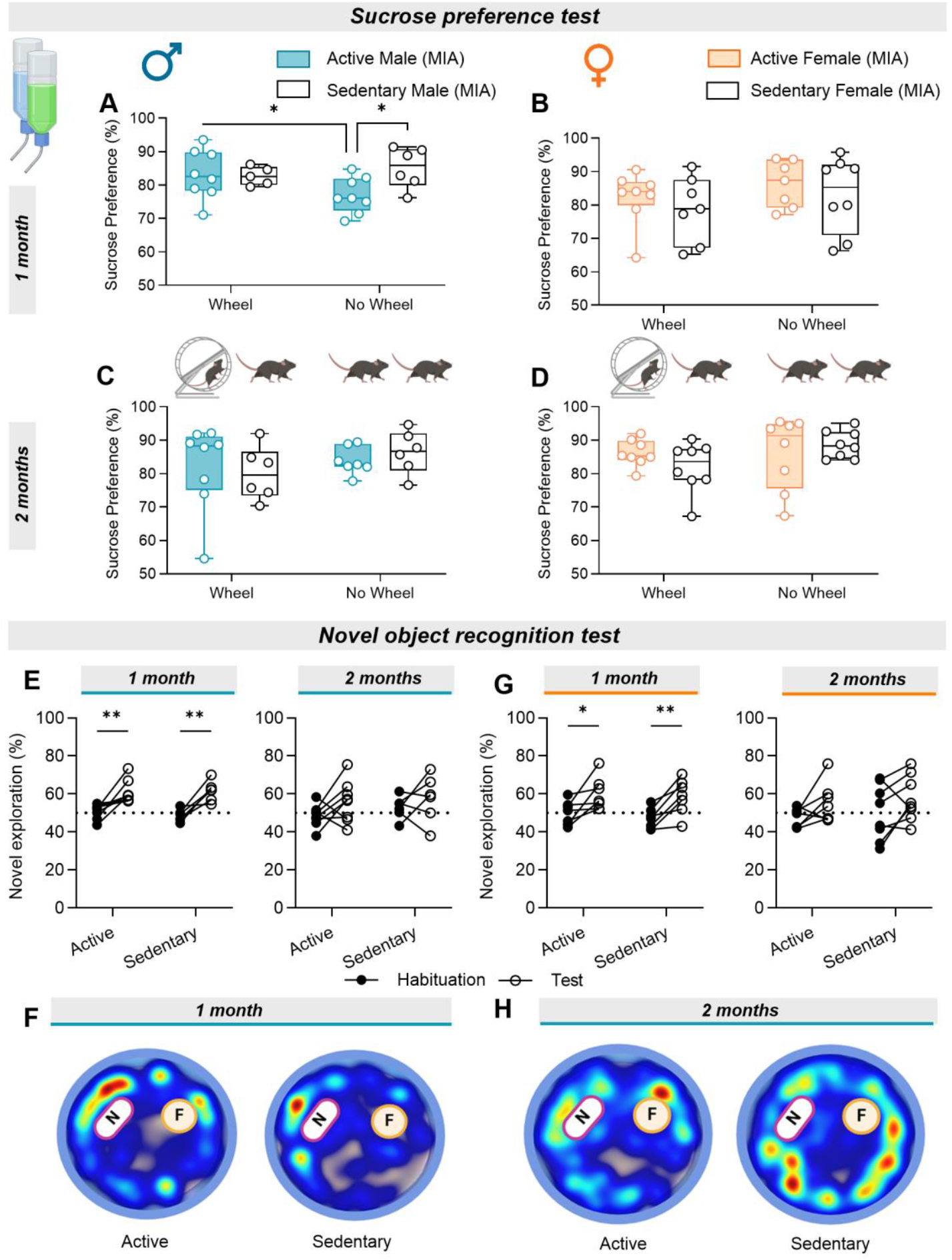
Wheel running does not change sucrose or novel object preference after the MIA injection. (**A**-**D**) Quantification of depressive-like behaviour using the sucrose preference test in the presence and absence of the running wheel at one month after the MIA injection in male (**A**) and female (**B**), and at two months after the MIA injection in males (**C**) and female (**D**) mice. Males: *N* = 5-8; Females *N* = 8/8. (**E-H**) Object novelty preference in male mice at one (**E**, left panel; **F**) and two (**E**, right panel; **H**) months after the MIA injection (*N* = 6-8), and in female mice at one (**G**, left panel) and two (**G**, right panel) months after the knee MIA injection (*N* = 7-8). Representative heat maps of mouse location during the NOR test (N – novel object, F – familiar object) in male mice at one (**F**) and two (**H**) months after the MIA injection.

Cognitive function was next assessed *via* the NOR task. At one month after the MIA injection, both male and female mice showed increased preference for the novel object, as evidenced by an increased amount of time spent exploring the novel object compared to the familiar one (***Fig. 3E-G***; ***E:*** left panel: 2-way RM ANOVA, ‘Time’ F_1,12_=30.55, *P* = 0.0001; ***G:*** left panel: 2-way RM ANOVA, ‘Time’ F_1,12_=23.45, *P* = 0.0004). This preference was no longer evident at 2 months, in either active or sedentary male and female mice (***Fig. 3***, right panels: ***E*** and ***G***; ***H***).

We next looked at anxiety-like behaviour. Using the open field test (***Fig. 4A-J***), no differences were observed between active and sedentary mice in the occupancy of the ‘Near Center’ zone of the arena and distance travelled within this zone at either one (***Fig. 4A-D, I***) or two months (***Fig. 4E-G, J***) post knee injury. Similar findings were obtained when assessing the probability of mice being located in the ‘Away from Center’ (***fig. S5A***) and ‘Far away from Center’ zones of the arena (***fig. S5B***), as well as locomotion (***fig. S5C, D***).

**Figure 4.**
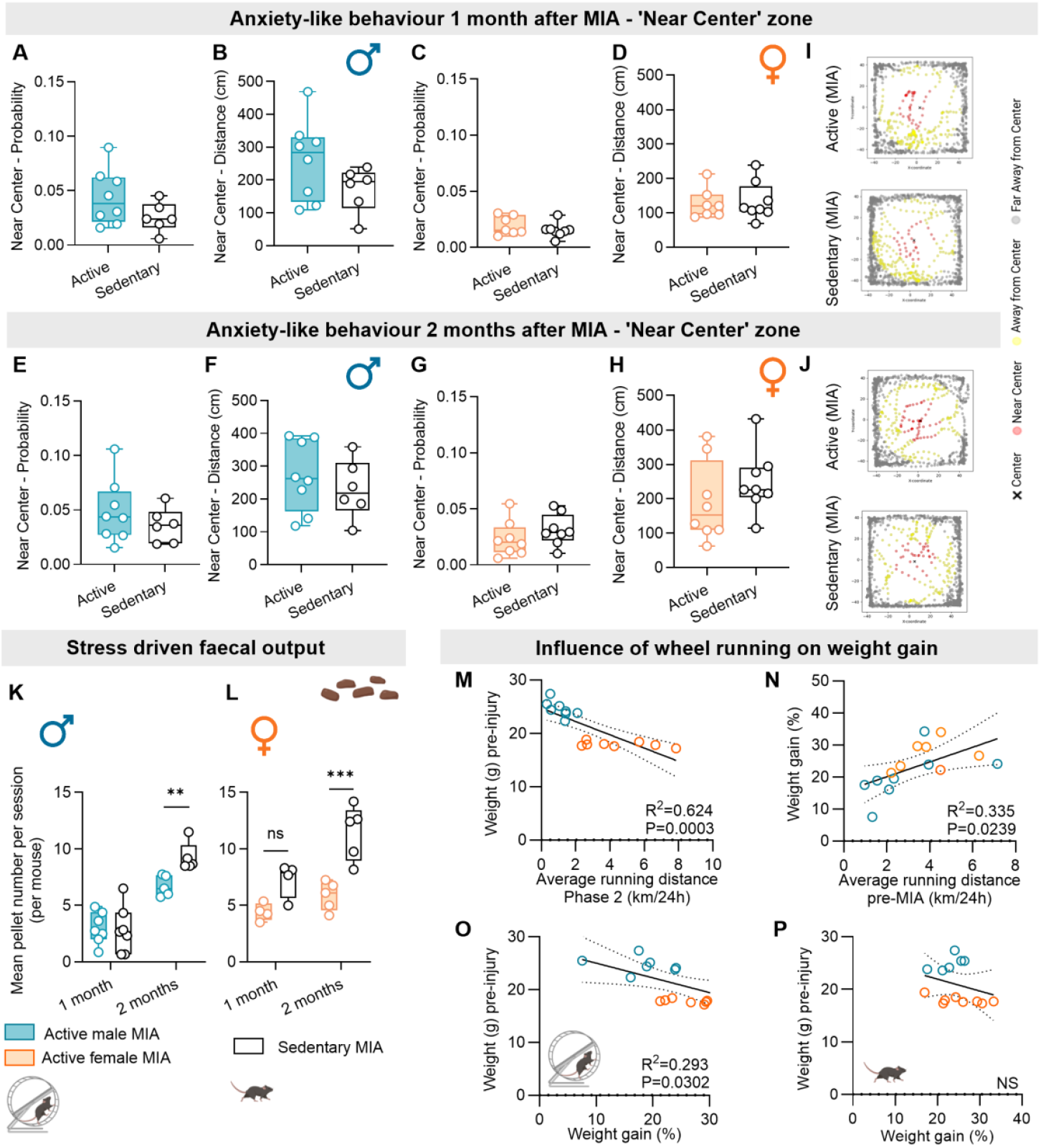
Wheel running does not change anxiety-like behaviour in the open field but attenuates stress-related responses measured as faecal pellets and influences weight gain. (**A**-**D**) Quantification of anxiety-like behaviours in the center of the open field arena at one month after the MIA injection in male (**A, B**, *N* = 8/6) and female (**C, D**, *N* = 7-8) mice. (**E**-**H**) Quantification of anxiety-like behaviours in the center of the open field arena at two months after the MIA injection in male (**E, F**, *N* = 8/6) and female (**G, H**, *N* = 8/8) mice. (**I, J**) Representative scatter plots illustrating the spatial distribution of the female mice movements across the defined zones of the open field arena at one (**I**) and two months (**J**) post-MIA injection. (**K, L**) Quantification of faecal pellet output during each session of von Frey testing sessions after MIA injection in male (**K**) and female (**L**) mice. Data points represent single sessions per animal. (**M**) Correlation between average running distance in Phase 2 and body weight pre-injury (MIA injection). (**N**) Correlation between average run before MIA injection and % weight gain at time of euthanasia (TOE). (**O**) Correlation between weight pre-injury and % weight gain at TOE in active mice. (**P**) Correlation between weight pre-injury and % weight gain at TOE in sedentary mice.

Stress is another factor that, when sustained, can have detrimental effects on both the pain and affective states [35]. To indirectly monitor the stress reactivity in our study, we quantified the faecal pellet output during mechanical sensitivity testing, as increased defecation is a well-established physiological outcome of acute stress. In both males and females, greater defecation was seen in sedentary mice compared with active mice at 2 months post-MIA (***Fig. 4K***, Mann-Whitney for non-parametric data, ‘Intervention’ F_1,20_=4.48, *P* = 0.0079; ***Fig. 4L***, 2-way RM ANOVA, ‘Intervention’: F_1,14_=25.54, *P* = 0.0001).

Lastly, while body weight measurements revealed no significant effect of wheel access in average weight or percent weight gain at the time of euthanasia in either sex (***fig. S6A, B***), there was a negative correlation between the running distance (Phase 2) and body weight pre-MIA injection (***Fig. 4M***) and at the end of the experiment (***fig. S6C***). There was also a positive correlation between percent weight gain and average running distance before injury (***Fig. 4N***), and a negative correlation between percent weight gain and weight pre-injury, in active mice only (***Fig. 4O, P***). Overall, the data point to baseline body weight as a potential key determinant of running behaviour, with lighter mice running more and displaying greater relative weight gain - a relationship that emerged only in wheel-access mice, consistent with the phenotypic relationship between body weight changes and physical exercise.

Collectively, these findings suggest that prolong voluntary physical activity does not significantly alter anxio-depressive-like behaviours or novelty preference during the chronic stages of MIA-induced osteoarthritic pathology. However, differences in the physiological responses to stress and body weight regulation were significant, indicating that exercise may exert more subtle influences.

## 4. Discussion

In this study, we report that voluntary wheel running, initiated prior to the induction of OA-like pathology and maintained for 2 months post-disease onset improved secondary mechanical hypersensitivity at the ipsilateral hindpaw but exacerbated some of the MIA-induced gait impairments. While activity had no obvious impact on the anxio-depressive-like behaviours and cognitive functioning, it reduced the physiological stress responses in both sexes, as assessed *via* faecal pellet output, and also impacted body weight gain. Together, our data suggest that voluntary physical activity in socially isolated mice can reduce hypersensitivity away from the joint while exacerbating gait deficit across sexes. These somewhat conflicting observations could explain the absence of obvious impact of voluntary wheel running on affective and cognitive outcomes.

Following the knee MIA injection, both male and female mice showed a decline in running activity, as previously observed in this animal model of OA [19,36]. This reduction in the acute post-injury phase was sex-dependent, with males exhibiting supressed activity for more than a week, while females returned to their pre-MIA activity levels within a few days. Interestingly, these sex differences persisted into the later stages of joint pathology. Male mice showed reduced daily running activity beginning around three weeks post-injection and continuing through the end of the study, whilst females sustained running levels comparable to their pre-injury baseline for 2 months. These findings matched our assessment of activity in home cages with non-invasive passive infrared recordings in naïve and MIA mice (***fig. S1*** and [27]).

Access to a running wheel improved the MIA-induced secondary mechanical hypersensitivity at the ipsilateral hindpaw, a hallmark feature of this OA model [27,37]. This improvement positively correlated with the distance run, suggesting a functional relationship between exercise and secondary pain relief. The positive outcome of running on secondary hypersensitivity did not extend to joint function, as evidenced by exacerbation of several MIA-induced gait impairments in active mice. These observations suggested that while physical activity had reduced secondary mechanical hypersensitivity, it may have exacerbated the joint pain, as seen by the diminished ground contact of the injured hindlimb during locomotion. The divergent effects of exercise on secondary mechanical hyperalgesia and joint pain were unexpected, given that our previous work demonstrated a strong positive correlation between these measures from joint disease onset through to three months [24,27]. This dissociation may indicate that distinct underlying mechanisms mediate exercise-induced modulation of central sensitization, including descending pain modulation, and joint-specific nociception, highlighting the need for future studies to disentangle these processes and their relevance for therapeutic interventions. An alternative explanation for the increased gait deficits is that it reflects an accelerated loss of joint structural integrity by exercise on an already compromised joint, rather than heightened pain per se. As previous studies using the MIA model of OA have shown that joint architecture undergoes rapid and extensive degeneration in both male and female mice [27,31,32], histological analysis of knee joint would need to be performed at the early stages of the disease to explore this possibility.

Our findings were also unexpected in the light of extensive literature supporting the beneficial effects of sustained exercise regimes on joint pathology [18,19]. However, previous findings in the MIA model revealed similarly complex outcomes, with voluntary exercise increasing bone remodelling in arthritic joints, suggesting that increased usage of a compromised joint can have negative consequences under certain conditions [15]. It has been proposed that moderate-intensity exercise regimes are effective therapeutic interventions for both joint pain and structural integrity, while intense exercise can exacerbate joint pathology [18,38]. While distances ran by mice given access to wheel are much greater than those covered in forced treadmill exercise, forced treadmill paradigms usually impose a sustained pace higher than the speed mice would voluntarily select, representing a more continuous physiological and mechanical load on the limbs [39–41]. Hence, it could be that the lower intensity of voluntary activity in our study may have been sufficient to attenuate secondary MIA-induced mechanical hypersensitivity, but inadequate to prevent the progression of OA pathology. This could explain why, while female mice ran far longer distances than male, we did not observe any significant sex-differences in pain-relief as both males and females running regimen would have been milder than forced exercise.

Given the strong relationship between sensory and affective aspects of pain, we also assessed the impact of exercise on anxio-depressive-like behaviours, known to develop in this model of OA at later stages of pathology [27,42]. We found no differences in the anxiety-like behaviours assessed using the open field test at one and two months after the MIA injection. This was unexpected given that previous studies have shown that regular physical activity reduces anxiety-like behaviours in rodents [43]. Similarly, no significant differences in sucrose preference were observed between active and sedentary mice, suggesting that wheel running does not influence depressive-like behaviours in single-housed mice with OA.

Nonetheless, exercise exerted a beneficial effect on physiological stress responses mounted during mechanical sensitivity testing. Male and female mice with access to a running wheel produced fewer faecal boli than their sedentary counterparts, indicating attenuated stress responses to repeated behavioural testing. This finding is consistent with previous work showing that exercising rats exhibited sustained reductions in blood corticosterone concentrations following intra-articular CFA compared with sedentary controls [44]. Interestingly, we found that lighter mice ran more and exhibited greater relative weight gain, an outcome not observed in sedentary mice and likely driven by increased energy expenditure and altered body composition (percent lean *vs* fat). These results indicate that voluntary running amplifies the relationship between early body mass and subsequent body weight.

While we report that most behavioural outcomes investigated in this study were mildly impacted by access to a running wheel, several limitations must be considered. First, the design of the study involved prolong social deprivation. Social isolation, particularly in a social species, is known to promote the emergence of anxio-depressive behaviours, impact reward-seeking and memory processes, and elevate basal glucocorticoid levels [45–47]. In addition, the outcome of the SPT used to evaluate depressive-like behaviour presents interpretation challenges in the context of our study. Research has shown that exercising mice often display reduced palatable food consumption [48,49], which may reflect substitution of reward or metabolic and energy balance adaptations, rather than true anhedonia. We selected the SPT specifically to minimize additional stressors and pain known to occur with alternative assays such as the forced swim or tail suspension test. While wheel running dimensions were optimised to fit within standard individually ventilated cages, their smaller diameter may have been less comfortable and more physically demanding for males to use as their body size increased [41], further limiting the interpretation of our findings. In the absence of a non-injured group, we could not determine whether MIA induced anxio-depressive-like behaviours in our cohorts, particularly given that these behaviours are known not to develop uniformly in mice with persistent pain [27].

In conclusion, our findings demonstrate that voluntary wheel running in the MIA mouse model of OA attenuates the mechanical hypersensitivity away from the joint while exacerbating gait impairments in both sexes. Although affective and cognitive outcomes remained unchanged, the reduced stress responses and impact on body weight in exercising mice suggest a degree of physiological benefits. Overall, these results indicate that while voluntary exercise might beneficially modulate central pain mechanisms, increased joint loading can compromise functional outcomes. Future work should aim to define exercise conditions that preserve the analgesic benefits while minimizing potential joint damage, thereby refining exercise-based therapies for OA pain management.

## Acknowledgments

This project was funded by a Versus Arthritis grant to SG, Research Award 21972. LA was funded by a grant from the Medical Research foundation MRF-087-0001-F-KOCH-C0917. We thank Prof Rob Brownstone, UCL, for giving us access to the wheel running set up used in Fig.S3.

## Authors Contributions

RF and SMG conceived and design the study and wrote the paper. RF ran all behavioural experiments. SH contributed to set up of the MIA model and some behavioural paradigms. SH and AL contributed to data analysis and interpretation. AW performed the immunofluorescence experiment, data acquisition, and analysis for fig. S3C, D. All authors reviewed and approved the final article.

## Conflict of interest

The authors have no conflict of interest.

## Supplementary Methods

### Methods

#### Animals and study design

A total of 16 male and 16 female C57Bl/6J mice were ordered from Charles River UK at 6 weeks of age. Upon arrival at the Biological Service Unit facility at UCL, they were single housed in individually ventilated cages containing sawdust bedding material, a wooden chewstick block, one cardboard tunnel, and Comfort Fresh nesting material, with the intervention cage also containing a vertical running wheel. Food and water were provided *ad libitum*, with animals being housed in an alternating 12/12h light/dark cycle (lights gradually on between 7-8am) and temperature and relative humidity-controlled conditions (22 ± 2ºC, 50 ± 10%). All procedures were carried out in accordance with the guidelines of the UK Animals (Scientific Procedures) Act 1989 and subsequent amendments.

A minimum of two week-period of habituation was allowed to ensure mice with access to the wheel would reach relatively stable levels of activity. After baseline measurements, mice received a unilateral MIA injection into the knee joint at ∼8-9 weeks of age. Two male mice had to be euthanised after the MIA injection leaving the final group sizes for males as follows: 8 mice in the active group (housed in cages with wheel) and 6 mice in the sedentary group (housed in cages without wheel). See ***fig. S1*** for a schematic illustration of the experimental timeline.

**Figure S1.**
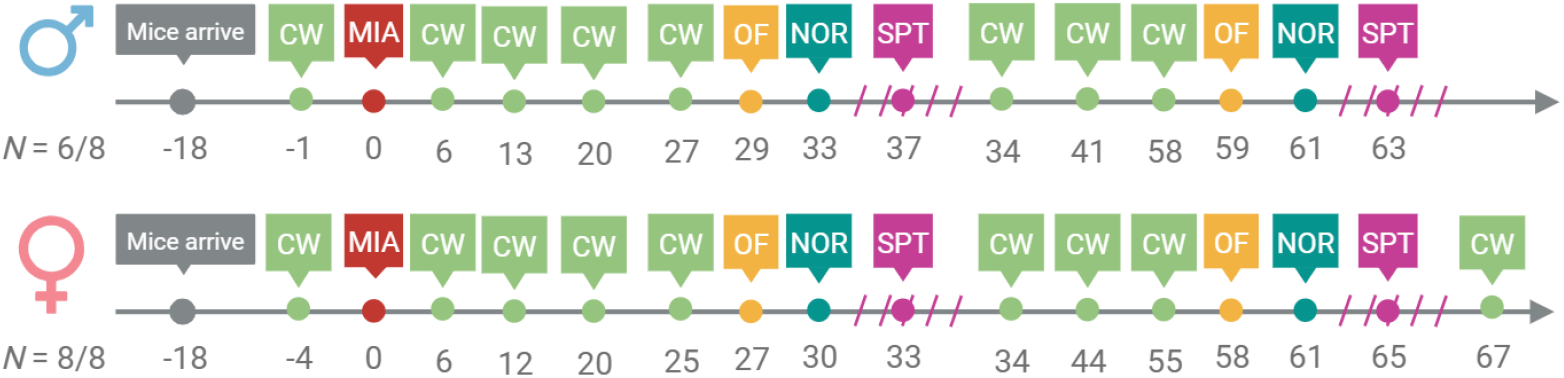
Timeline of experimental procedures and behavioural tests carried out. CW – CatWalk; MIA – monoiodoacetate; OF – open field test; NOR – novel object recognition test. Slashed lines around SPT indicate that the test was conducted over several days.

#### Activity recording

Running activity was quantified using an activity wheel monitoring system (Columbus Instruments, Columbus, OH, USA). Intervention cages were equipped with a vertical polycarbonate running wheel (10.5cm diameter) and wheel revolutions were automatically detected using a Hall-effect sensor connected to an eight-channel counter and Quad CI-Bus interface. Data were recorded using Multi-Device Interface (MDI) software (version 1.11.7). Running activity was continuously monitored over a 24h period. Cages were checked regularly to ensure unobstructed access to the wheel. Data were excluded from analysis if wheel movement was impeded (*e*.*g*. by cardboard tunnel or sawdust) or if hardware or software malfunctions occurred.

#### Monoiodoacetate-induced arthritis

To induce osteoarthritis, mice received an intra-articular injection of monoiodoacetate (MIA) in the left knee joint as previously reported [1,2]. On the day of injection, mice were placed under general anaesthesia using a gas mixture of isoflurane in O_2_ (2.5% for induction, 2% for maintenance, 1.5L/min flow rate). After cessation of reflexes, the area surrounding the knee joint was trimmed and disinfected. The knee was held in a 90° bend position and 10μl of 10% w/v MIA solution (Sigma-Aldrich, I2512) was injected through the patellar tendon in the synovial space using a micro-fine 0.3ml 30G insulin syringe (BD, BD3/A1). After needle retraction, the hindleg was gently moved back-and-forth to ensure even distribution of the solution. Mice were then returned to their home cages and monitored closely for the next 72h, time during which no behavioural measurements were taken.

#### Mechanical sensitivity

Hindpaw mechanical sensitivity was assessed using the up-and-down method as previously described [2–4]. Mice were individually placed in plexiglass cubicles (9.5cm x 9.5cm, 14cm height) on an elevated wire grid (Ugo Basile) and allowed to habituate to the room and experimenter for 30-60min before testing. Calibrated von Frey monofilaments (Ugo Basile, Italy) were applied to the plantar surface of the hindpaw to determine the 50% withdrawal threshold: starting with a monofilament of 0.6g, a positive response involving a withdrawal response would be followed by the application of the next, lower force filament (0.16g). Conversely, the next higher force filament would be applied (1g). After a first change in pattern, four additional stimulations were applied and the 50% paw withdrawal threshold was calculated using the formula: 50% *mechanical threshold* (log(*g*)) = 10^log(last filament weight (g))+*k*\*0.3^, where *k* is a constant value corresponding to each individual response pattern obtained [3].

#### CatWalk gait analysis

Gait behaviour was analysed using the CatWalk^®^ XT 10.0 system (Noldus Information Technology), as we previously described [2]. The apparatus consists of a suspended glass runway internally illuminated by green LED light. Two parallel panels create a corridor which the mouse has to transverse, while a red LED illuminated ceiling offers contrast for optimal detection of paw prints by a high-speed camera positioned underneath the glass plate. Mice were placed at the beginning of the runway and allowed to freely explore the setup with minimal intervention form the experimenter. No habituation to the room or the setup was provided to reduce the reduced likelihood of mice to explore the space. A compliant run was considered when the mouse walked along the corridor at a regular speed, without slowing down, pausing, or turning around. The green intensity threshold (0.10) and camera gain (19.58) were constant throughout the experiments for inter-animal paw comparison. A minimum of two compliant runs were included in the analysis, while animals that were reluctant to run on specific days were excluded from analysis. Outcomes are expressed as a ratio between the ipsilateral (left) and contralateral (right) hindpaws. The parameters analysed were based on previous literature on arthritis models [5].

#### Open field test

Anxiety-like behaviour was assessed using a squared open field arena (90cm length, 40cm height). Mice were allowed a 40min habituation to the room, before being placed individually in the center of the open field arena. Subject’s movements were live tracked and recoded for 5min using the Ethovision XT14 software (Noldus Information Technology). The arena was cleaned with 70% ethanol after each trial.

Raw movement data (x-y coordinates and movement velocity) were exported and analysed using a Python script. The geometric center of the arena was determined as the midpoint between the minimum and maximum x and y coordinates recorded during each trial. Using this midpoint, three concentric circular zones were defined: ‘Near Center’ zone (within a 20cm radius, R ≤20cm), ‘Away from Center’ zone (20cm ≤ R ≤ 40cm), and ‘Far Away from Center’ zone (40 ≤ R ≤ 60cm). For each frame, the Euclidean distance between the animal’s position and the arena midpoint was calculated to classify the position into one of these zones.

Traveling velocity values, exported from EthoVision, were averaged for each defined zone to indicate the animal’s movement pattern. Total distance moved in each zone was calculated by summing the distance moved every 0.2sec within the three zones. To determine the probability of the mouse being located in a specific zone, a Boolean array mask was created where each element was either True (1) if the mouse center point fell within the respective zone, or False (0) otherwise. The mean value of each mask array represented the probability for each respective zone. Representative images were created using the scatter function from Matplotlib to plot mouse center points that fall within each mask, using different colours for each zone.

#### Sucrose preference test

Anhedonia was assessed using the sucrose preference test as previously described [2]. One week before testing, mice were habituated in their home cages to drinking water from two 50ml falcon tubes fitted with a nozzle. Three days before testing, the water in one of the two bottles was changed to a 1% sucrose solution, and presented to mice for 12h, from 7pm to 7am the next day. In the following night, the position of the sucrose solution relative to water was swapped, to ensure mice would become familiar with the palatable solution being present on either side. No access to sucrose solution was provided in the 24h time period before test, while no food or water deprivation was employed throughout the study [6].

On the first night of testing, the bottle containing the 1% sucrose solution was placed in the food hopper on the left side relative to the water bottle; on the second night, it was placed on the right side to control for potential side preferences. Bottles were weighed at the start (7pm) and end (7am next day) of each testing period, and sucrose preference was calculated using the formula: 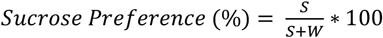 where *S* is the volume of sucrose consumed, while *W* is the volume of water consumed overnight. To account for possible competing rewarding effects between the running activity and sucrose consumption, testing was carried out over four consecutive nights. In the first two nights, mice had access to the running wheel (intervention group), and sucrose bottle was altered between left and right positions relative to water. On the final two nights, the test was repeated in the absence of wheel access for the same mice. No differences in sucrose preference were observed based on bottle position, hence only data from trials in which the sucrose bottle was placed on the right are presented here.

#### Novel object recognition test

To quantify novelty preference, the novel object recognition (NOR) test was employed as previously described [2,7]. After a 30min habituation period to the room, mice were individually placed, wall-facing in the far-end of a round arena (diameter 35cm, hight 30cm) containing two identical objects (circular glass bottles of diameter 6.5cm, height 17cm, and brown colour). A video camera positioned above the arena and connected to a computer tracked and recorded the mouse movements using the Ethovision XT14 software (Noldus Information Technology). This phase lasted 10min and it represents the ‘habituation’ phase. Three hours later, the mouse was reintroduced to the arena, in which this time around one of the familiar objects was replaced with a novel one (elliptical shaped glass bottle of length 8.5cm, width 4.5, height 15cm, and white colour). This phase lasted 5min and it represents the ‘test’ phase of the NOR test. During this phase, the location of the novel object in the arena was swapped relative to the familiar object in a random manner across trials to account for potential side preferences.

A mouse was considered to explore an object when its nose point was within a 2cm distance from one of the two objects present in the arena. The amount of time spent exploring the objects was calculated using the formula: 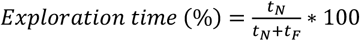, where *t*_*N*_ and *t*_*F*_ is the time spent exploring the novel and familiar object, respectively. For the habituation phase, the ‘novel’ object was treated as the familiar one that would later be replaced with the novel object during the testing phase.

#### Tissue collection and immunofluorescence

Mice were transcardially perfused with heparinised saline solution followed by 10% formalin (Sigma-Aldrich, HT-501640-19L). Brain tissue was removed and postfixed for an additional 2h then stored in 30% sucrose solution containing 0.5mg/ml NaN_3_ at 4°C. Hippocampal sections of 40μm thickness were washed with 0.1M phosphate buffer (PB), then incubated for 1h with 30% donkey serum (Biosynth, 88R-D001) in 0.1M PB containing 0.3% Triton X-100 at room temperature. Sections were then incubated overnight with rabbit anti-doublecortin (DCX, 1:5000, Abcam, ab18723) at 18°C. Next day, sections were washed in 0.1M PB then incubated for 2h with rabbit biotinylated secondary antibody (1:400; Bethyl Labs, A120-108B), followed by incubation in preformed avidin-biotin-HRP complexes (1:200; Vectastain ABCKit Peroxidase, PK-6100) for 30min. Finally, tissue was incubated for 5min with tyramide solution (ApeBio, A8011-APE), washed 2×15min with 0.1M PB, then incubated for an additional 2h with Avidin D, Fluorescein labelled (Vector Labs, A-2001-5). Sections were given a final wash, mounted on gelatinised slides, and coverslipped with Fluoromount Aqueous Mounting Medium (Sigma-Aldrich, F4680). DCX immunopositive cells were visualised and manually counted within the dentate gyrus of the hippocampus using a Leica (Nussloch, Germany) DMR microscope connected to a Hamamatsu (C4742-95; Shizuoka, Japan) digital CCD camera and Velocity 6.3 Software.

#### Statistics

Data were analysed using GraphPad Prism (version 10.5.0 (774)) and IBM SPSS statistics (version 331.0.0.0(117)). Individual animals were considered experimental units. Statistical analysis was mostly performed using Student’s *t* test, 1-way or 2-way repeated measures (RM) ANOVA, as well as simple linear regression. Normality was assessed using the Shapiro–Wilk test and inspection of Q–Q plots, which provide a more robust assessment when formal tests are overly sensitive. When sphericity or multivariate tests could not be computed due to an unbalanced data structure (*i*.*e*., an insufficient number of complete cases relative to the number of time points), we analysed the data using linear mixed-effects models, which do not rely on the sphericity assumption and are robust to missing or uneven repeated-measures data. For von Frey thresholds, raw data were log-transformed to normalize distributions and stabilize variance. This is a well-accepted transformation of von Frey data [8]. Residuals rarely deviated from normality and Q–Q plots showed no severe departures, supporting the validity of parametric analyses. Where deviations from normality occurred (only in isolated subgroups), corresponding non-parametric tests (Mann–Whitney’s test) was used. Significance was set at *P* < 0.05. Outcomes *t*, F, and *P* values are reported in the main text, while the *N* values (sample sizes) are provided in the figure legends. Mixed effects analysis was used for data-sets with missing data-points due to random factors, like software malfunctions or lack of compliance from the mice on the catwalk. Multiple comparisons were performed using Šídák’s method. When the assumption of sphericity was violated in RM ANOVA, the Greenhouse Geisser correction was applied. Data are presented as group means ± standard error of the mean (SEM), with the exception of box-and-whiskers plots, which display the IQR. Outliers were identified and excluded using Tukey’s method, based on IQR. Overnight wheel running activity was averaged over 3-5 days to reduce day-to-day variability and minimise the impact of data lacking due to unexpected software malfunction.

**Table S1.**
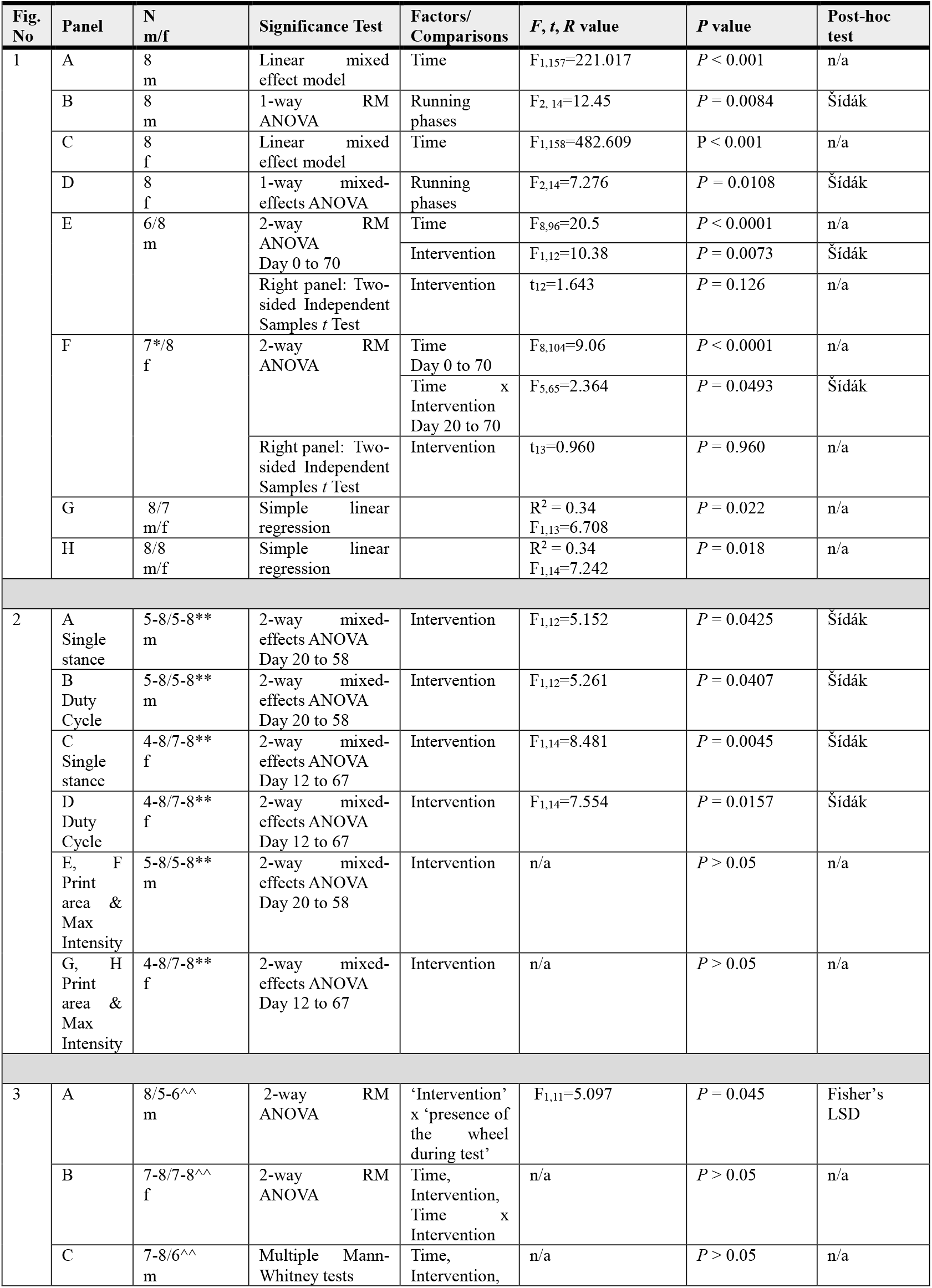
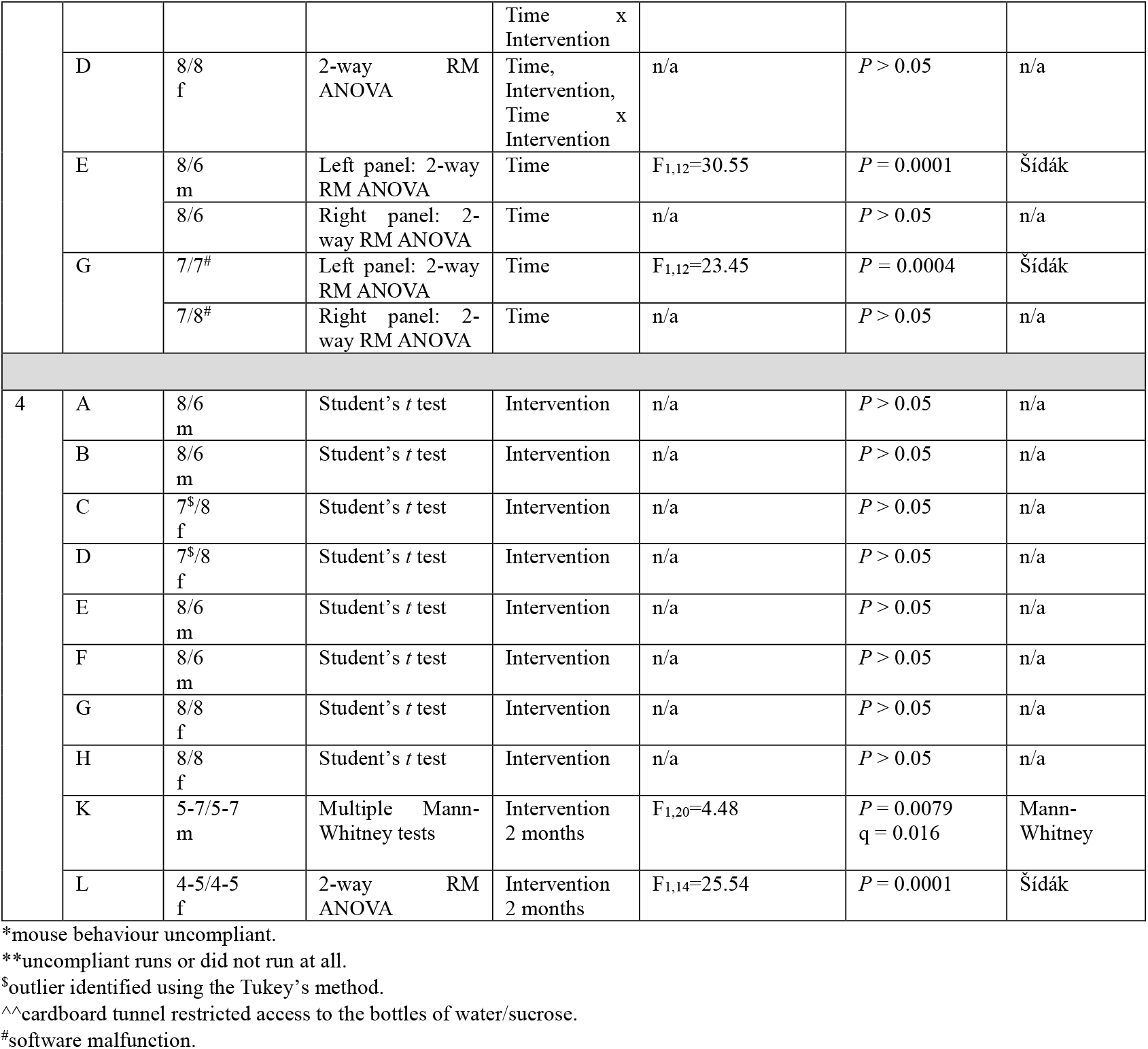
Description of statistical results for Figures 1 to 5.

## Supplementary Data

**Figure S2.**
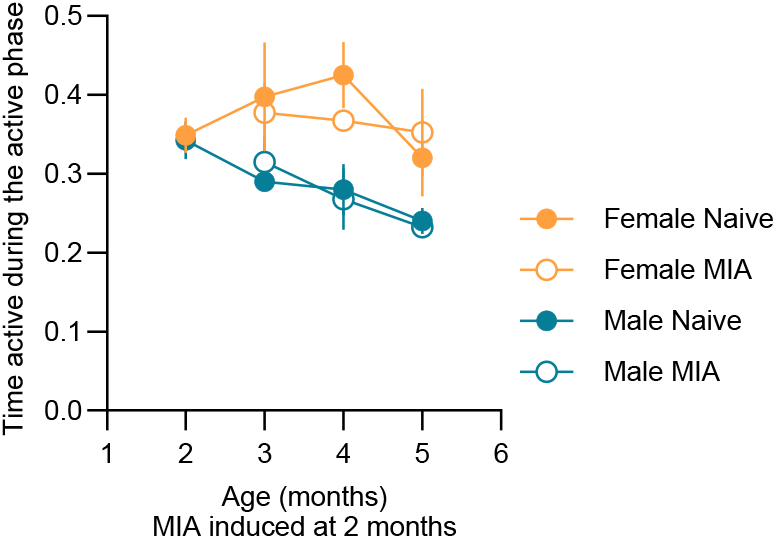
The homecage activity of male, but not female mice, decreases with age, regardless of injury. Activity was measured using non-invasive passive infrared sensors [9,10]. Time active is expressed as a ratio of total time. MIA was injected into the knee when mice were 2-month-old [2]. 3-way RM ANOVA 3 months – 6 months: ‘Time’: F_2,24_=4.205, *P* = 0.027; ‘Sex’: F_1,12_ = 12.606, *P* = 0.004.

**Figure S3.**
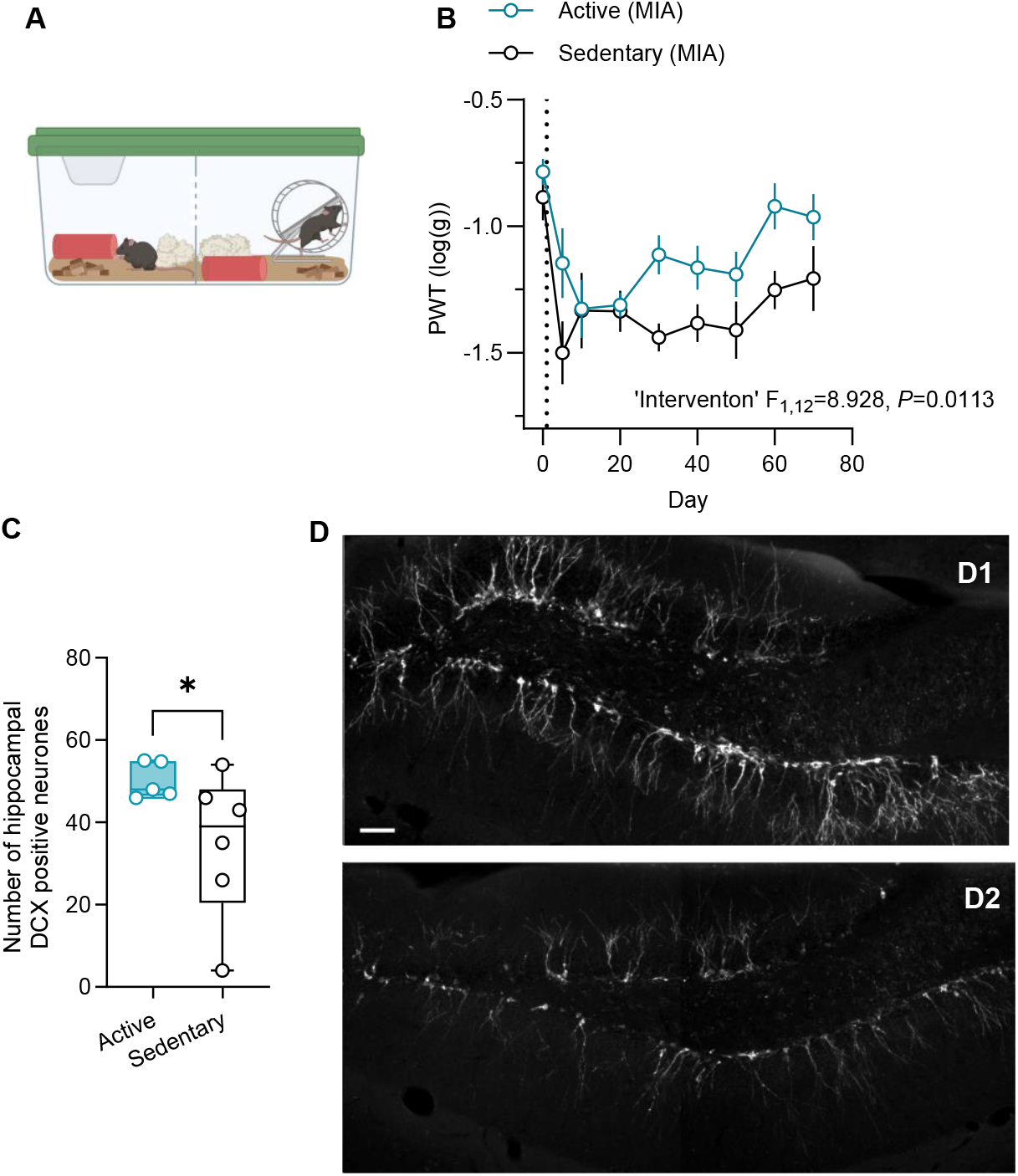
Wheel running attenuates MIA-induced mechanical allodynia and promotes neurogenesis in the hippocampus. **(A)** Schematic representation of the experimental setup in which two mice were housed in a single cage separated by a perforated plexiglass divider, allowing sensory (visual, olfactory and audio), but not physical contact between the two mice. **(B)** Hindpaw mechanical threshold at the ipsilateral hindpaw following the MIA injection (2-way RM ANOVA D0-70 ‘Intervention’ F_1,12_=8.93, *P* = 0.0113; N = 7/7, male mice). (**C**) Quantification of hippocampal doublecortin (DCX) expression at 2 months post MIA injection, expressed as the sum of the ipsilateral and contralateral sides of the dentate gyrus (Welch’s *t*-test. t=2.057; df=5.715; *P* = 0.0439). (**D**) Typical DCX immunostaining in the hippocampal dentate gyrus of active (D1) and sedentary (D2) mice. Scale bar = 50μm.

**Figure S4.**
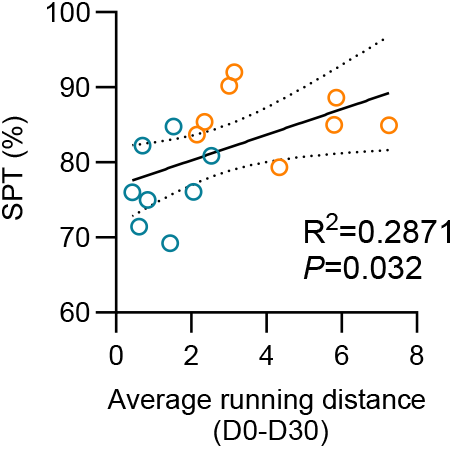
In the absence of a wheel, there is a positive correlation between sucrose preference at one month and running in active mice. Correlation between mean distance ran between day 0 to day 30 (km/24h) and sucrose preference (%) at one month after the MIA injection.

**Figure S5.**
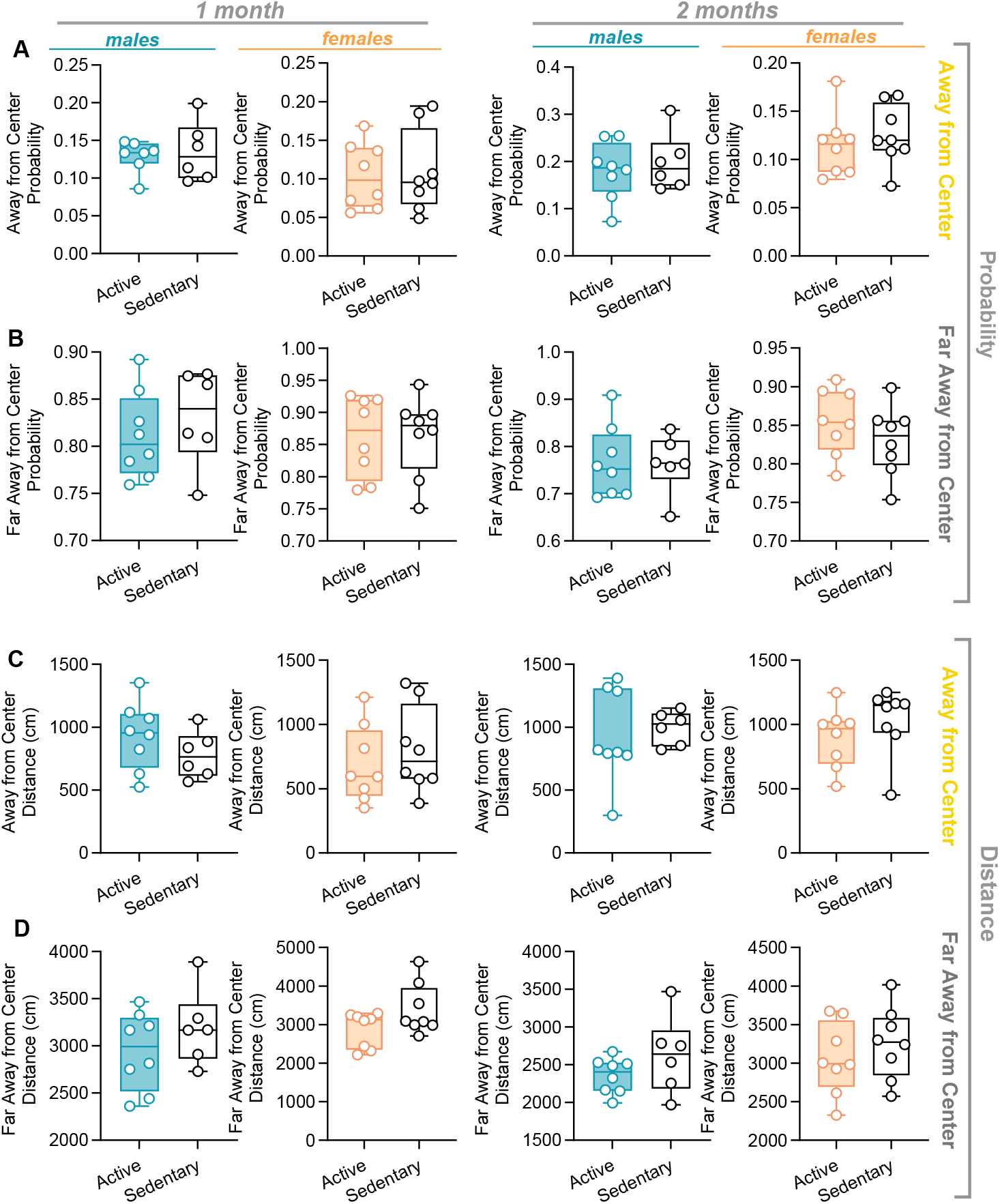
Access to a running wheel does not change anxiety-like behaviours assessed using the open field test in mice. (**A-D**) Quantification of anxiety-like behaviours in the ‘Away from Center’ (A, C) and ‘Far away from Center’ (B, D) zones of the open field arena in male (N = 6-8) and female (N = 8/8) mice at one and two months after the knee MIA injection.

**Figure S6.**
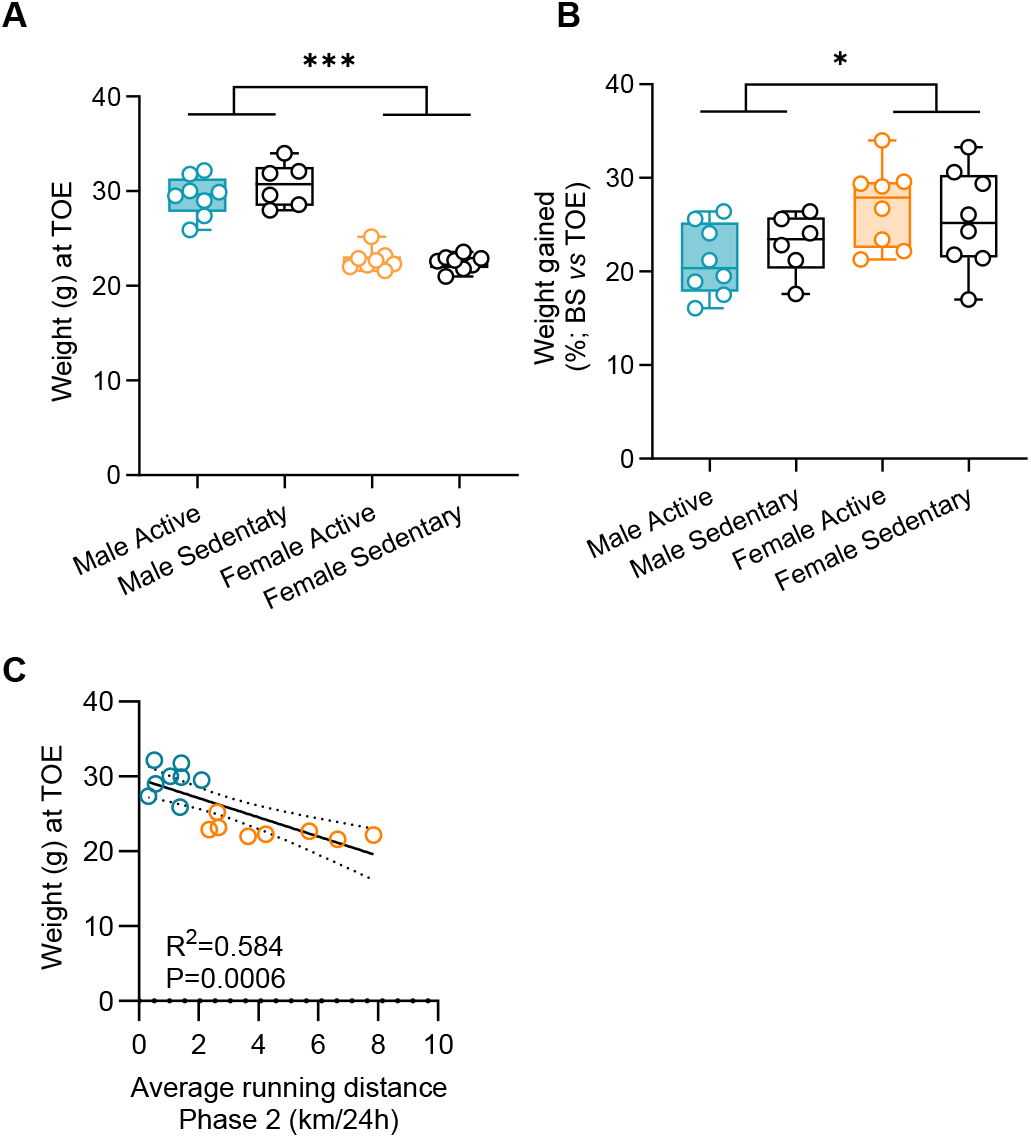
There is a negative correlation between the running distance and body weight of mice at the end of the experiment. (**A)** Body weight (g) at time of euthanasia (TOE); 2-way ANOVA ‘sex’: F_1,26_=150.78, *P* < 0.001. *N*= 6-8. **(B**) Percent body weight gain from baseline (BS; prior to knee MIA injection) to the end of the experiment (TOE); 2-way ANOVA ‘sex’: F_1,26_=6.731, *P=*0.015. *N* = 6-8. (**C**) Correlation between average running distance in Phase 2 and body weight at TOE.

## Notes

### Competing Interest Statement

The authors have declared no competing interest.

